# Orbitofrontal circuits for context-gated reward predictions

**DOI:** 10.64898/2026.03.05.709962

**Authors:** Sophie Peterson, Margo Le, Neil Dundon, Ronald Keiflin

## Abstract

The orbitofrontal cortex (OFC) is essential for the contextual regulation of reward predictions, yet the circuit mechanisms underlying this function remain unclear. Here, we dissected the roles of two major OFC efferent pathways —projections to the central dorsal striatum (CDS) and the mediodorsal thalamus (MDT)— in contextual regulation. Male and female rats performed a context-dependent discrimination task in which the reward-predictive status of two auditory target cues (X and Y) was determined by a visual contextual cue (A). Using temporally precise optogenetic inhibition of OFC terminals, we found that silencing OFC→CDS profoundly disrupted contextual gating, virtually abolishing the negatively gated component (A:Y− / Y+) while producing modest effects on the positively gated component (A:X+ / X−). In contrast, OFC→MDT inhibition produced modest and qualitatively distinct effects, characterized by a generalized increase in responding in context A and a decrease in context noA. These effects were reproduced in a connectionist model in which context influenced predictions either additively (via elemental representations) or by acting as a gatekeeper (via configural cue/context representations). Disrupting the configural pathway mimicked OFC→CDS silencing, whereas amplifying additive contextual inputs replicated OFC→MDT inhibition. Together, these findings reveal complementary circuit mechanisms through which the OFC shapes context-appropriate reward seeking.

## INTRODUCTION

Reward-predictive cues exert powerful control over motivated behavior. However, in naturalistic settings, reward cues are often ambiguous —what is rewarded in one context might not be rewarded in another context. To resolve this ambiguity, animals use contextual cues as hierarchical modulators —or gatekeepers— that guide the interpretation of ambiguous cues by controlling the retrieval of appropriate cue-outcome memories (e.g., the word “apple” evokes different expectations at the farmers market vs. the electronic store). This contextual gating of associative memories is a cornerstone of cognitive control and behavioral flexibility^1–7^.

Impairments in contextual gating have been implicated in several neuropsychiatric disorders —including obsessive-compulsive disorder and substance use disorders— contributing to the intrusive thoughts and maladaptive, context-inappropriate behaviors that characterize these disorders^3,4,8–11^. Yet, compared to our growing understanding of cue-elicited motivated behaviors, the neural circuits underlying the contextual gating —or regulation— of those behaviors remains largely unknown.

The orbitofrontal cortex (OFC) plays a crucial role in forming and using “cognitive maps”, i.e. internal models of the predictive relationships between events —including context-dependent relationships^12–15^. These OFC predictive maps can guide decision-making, particularly in complex situations where the interpretation of ambiguous cues requires integration of contextual information and knowledge of broader task structure. Consistent with this view, recent studies have implicated the OFC in context-gated reward predictions and the contextual regulation of reward seeking^16–20^.

While this prior work identified the OFC as a critical region for the contextual gating of reward predictions, the broader neural circuits involved in the contextual regulation of reward seeking remain unclear. OFC neurons project to multiple cortical and subcortical regions, with the central dorsal striatum (CDS) and mediodorsal thalamus (MDT) receiving particularly dense afferent inputs from the OFC^21–25^. Both of these regions have been implicated in cue-evoked motivated behaviors and executive control^26–33^. In light of these anatomic and functional characteristics, OFC projections to the CDS and MDT are strong candidates for mediating the context-informed, top-down regulation of cue-driven reward seeking.

The goal of this study was to examine the contribution of these specific OFC output pathways (OFC→CDS and OFC→MDT) in the contextual regulation of cued reward-seeking behavior. To that end, we trained rats in a context-dependent discrimination task in which efficient reward seeking behavior critically relies on the contextual gating of cued reward predictions (the context informing the predictive status of ambiguous reward cues). We then interrogated the functional role of these OFC subcortical output pathways by optogenetically silencing OFC terminals in either the CDS or the MDT. We found that the two OFC output pathways make distinct contributions to the contextual regulation of motivated behavior, revealing complementary circuit mechanisms through which the OFC shapes context-appropriate reward seeking.

## RESULTS

Male and female rats (n = 20; 10M + 10F) were trained on a context-dependent discrimination task (adapted from ^16,17^). Two auditory target cues (X and Y; 10s) were presented either in presence or absence of a sustained visual contextual cue (A; 2min) —the visual contextual cue informing the predictive value of the target cues. Specifically, cue X was rewarded with sucrose only in the presence of A, while cue Y was rewarded only in its absence (A:X+ / X− / A:Y− / Y+). Critically, this discrimination cannot be solved through simple binary cue–outcome, or context-outcome associations; instead, accurate performance relies on hierarchical, context-gated associative structures to interpret ambiguous cues and generate appropriate reward predictions (**Fig. 1A**).

**Fig. 1.**
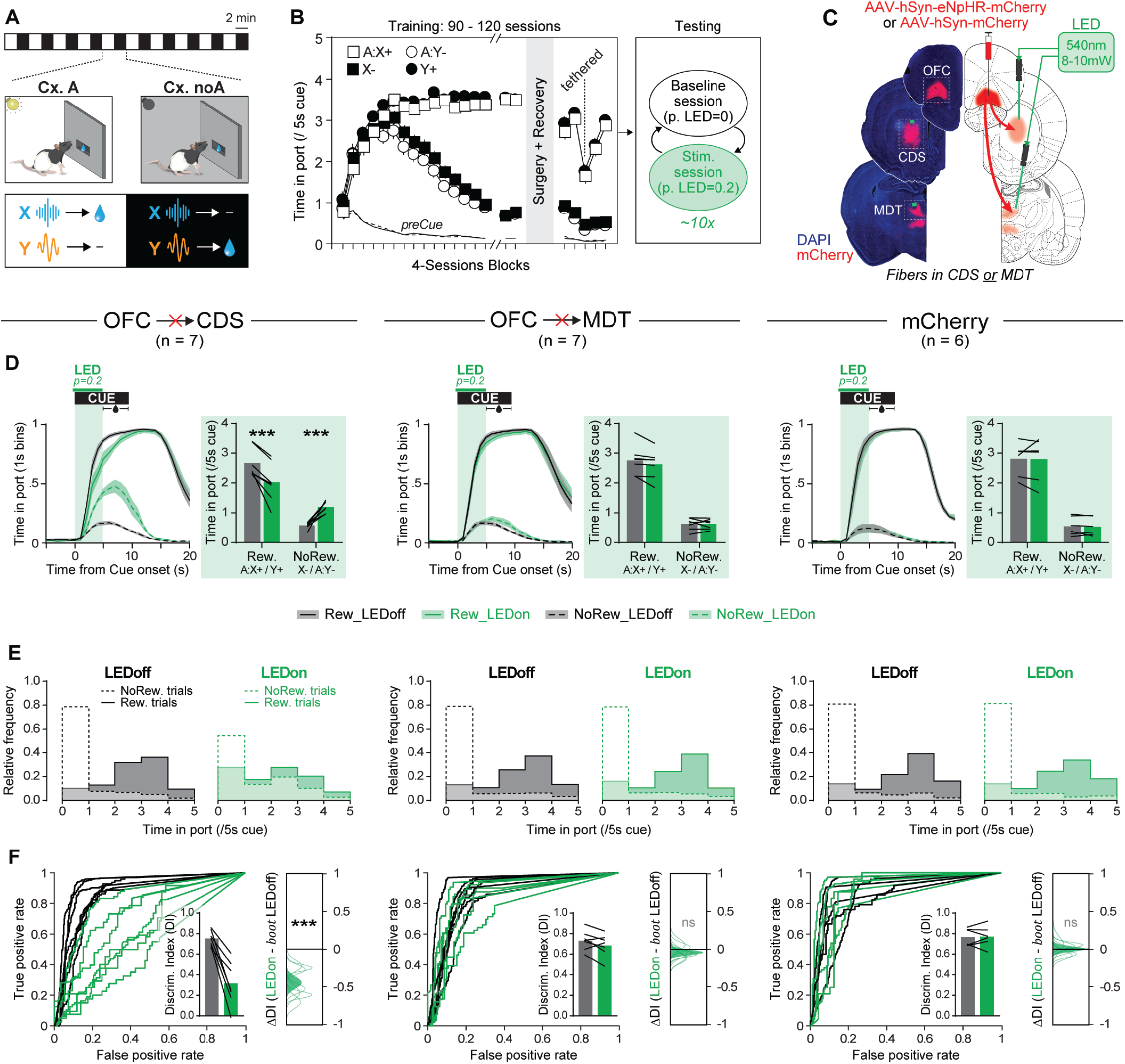
Phasic inhibition of OFC→CDS pathway profoundly disrupts the contextual regulation of cued reward predictions. **A**. Context-dependent discrimination task. Two auditory target cues (X and Y; 10s) were presented either in presence or absence of a sustained visual contextual cue (A; 2min). Cue–reward contingencies differed across contexts —the visual context informing the predictive value of each target cue. **B**. Experimental timeline. Following initial training, rats underwent surgery and post-surgical retraining. During subsequent stimulation sessions, LED stimulation was applied on 20% of trials. Stimulation sessions alternated with baseline sessions during which no stimulation was delivered. **C**. Right: Experimental strategy for silencing OFC→CDS and OFC→MDT pathways. Left: representative fluorescence images of (eN-pHR) mCherry viral expression in ventral OFC neurons and their axon terminals in the CDS or MDT, along with optical fiber placements in those target regions. **D**. Port occupancy throughout cue presentation. Histograms summarize mean port occupancy during the first 5 s of the cue (before reward delivery); lines represent individual subjects. OFC→CDS pathway silencing increased responding on non-rewarded trials and decreased responding on rewarded trials (*** P < 0.001, LEDoff vs. LEDon, LMM post-hoc). **E**. Distribution of port-occupancy durations. Under control conditions (LEDoff), the distributions for rewarded and non-rewarded trials were clearly separated; this separation was reduced during OFC→CDS silencing (LEDon). **F**. ROC analysis of behavioral discrimination. Individual-subject ROC curves were constructed from responses to rewarded (true positive) and non-rewarded (true negative) trials. Histogram shows the average discrimination index (DI) computed from the area under the ROC curve (DI = 2 × [auROC − 0.5]). Insets: effect of LED stimulation on discrimination performance expressed as ΔDI (LEDon − bootstrapped LEDoff). Distributions show bootstrapped ΔDI estimates for individual subjects (green lines) and the group-level estimate (green area). OFC→cDS pathway silencing significantly impaired discrimination performance (*** P < 0.001, LEDoff vs. LEDon, bootstrap test). LED stimulation appeared to have no effect on cue-evoked responding or discrimination performance in the OFC→MDT group or in m-Cherry control animals.

Following acquisition of discriminated (i.e. context-sensitive) Pavlovian approach (∼100 sessions), rats were divided into subgroups of matched performance and received injections of AAVs for the expression of inhibitory opsin eNpHR, or the control protein mCherry, in the ventrolateral OFC. Bilateral optical fibers were then implanted, aimed at either the CDS or the MDT, allowing for the optogenetic silencing of OFC axon terminals in these target regions (**Figs. 1B-C and S1**). Light delivery in the CDS or MDT had no detectable effect in rats lacking the inhibitory opsin; therefore, to minimize animal use, all these rats were pooled into a single mCherry control group. This resulted in three experimental groups: NpHR: OFC→CDS (n = 7), NpHR: OFC→MDT (n = 7), and mCherry control (n = 6).

To interrogate the functional role of the targeted pathways in the contextual regulation of cued reward predictions, LED light (540nm; 8-10mW) was delivered to the CDS or MDT for 5s, starting 200ms before cue onset; this time window corresponding to the anticipatory reward-seeking period. During optogenetic inhibition sessions, LED light was delivered semi-randomly on 20% of trials of each type (A:X+ / X− / A:Y− / Y+). These stimulation sessions alternated with baseline sessions without optogenetic manipulation, in order to minimize potential stimulation-induced neuroadaptations and assess the effect of optogenetic manipulations against stable baseline performance. Importantly, performance on LEDoff trials within stimulation sessions did not differ from performance during baseline sessions, ruling out within-session carryover effects of LED stimulation and confirming that optogenetic effects were temporally specific and restricted to stimulated trials (**Fig. S2**). All results reported below compare LEDon vs LEDoff trials within stimulation sessions.

### Phasic inhibition of OFC→cDS pathway profoundly disrupts the contextual regulation of cued reward predictions

Cue-evoked reward prediction was assessed by measuring the time rats spent in the reward port (i.e. port occupancy) during the first half of the cue period, prior to reward delivery—reflecting anticipatory reward-seeking behavior. By the end of training —prior to optogenetic silencing— all rats exhibited significantly greater responding on rewarded trials (A:X+ and Y+) compared to nonrewarded trials (X− and A:Y−), indicating robust contextual modulation of reward prediction (OFC→CDS: Discrimination Index (DI) = 0.738; OFC→MDT: DI= 0.730; mCherry: DI = 0.786; P = 0.460).

To assess the effect of LED stimulation, we first fit a linear mixed model (LMM) with Trial Type (rewarded vs. nonrewarded) and LED condition (on vs. off) as within-subject factors, and Group as a between-subject factor. This omnibus analysis revealed a significant Trial Type × LED × Group interaction (F(2, 17534.1) = 72.6183, p < 0.001), indicating that the impact of LED stimulation varied across groups and trial types. To further characterize the effects of LED stimulation, separate LMMs were then fit within each experimental group. This analysis revealed that inhibition of the OFC→CDS pathway disrupted anticipatory reward seeking in a trial-dependent manner (Trial Type × LED interaction: F(1, 5746.1) = 237.9219; P < 0.001). Specifically, LED stimulation increased responding to nonrewarded cues (X− and A:Y− combined) and decreased responding to rewarded cues (A:X+ and Y+ combined) (pairwise comparisons of estimated marginal means, Ps < 0.001). This pattern indicates a clear impairment in the contextual regulation of reward prediction following OFC→CDS pathway inhibition. In contrast, inhibition of the OFC→MDT pathway or LED light delivery in control rats lacking eNpHR, had no significant effect on cue-evoked reward seeking (main effect of LED: Ps>0.2069; Trial Type × LED interaction: Ps > 0.1289) (**Fig. 1D-E**).

To facilitate group comparisons, we calculated for each rat, a Discrimination Index (DI) based on the area under the receiver operating characteristic curve (auROC) formed by the distributions of responding to rewarded versus nonrewarded cues. The DI was computed as DI = 2*(auROC − 0.5), thereby rescaling the auROC to range from −1 to 1, where 1 indicates perfect discrimination (responses to rewarded cues always exceeding responses to nonrewarded cues), and 0 indicates no discrimination. Separate DIs were calculated for trials with and without LED stimulation (DI_LEDon and DI_LEDoff), allowing us to quantify the effect of LED stimulation on discrimination performance. Within-subject comparisons confirmed that inhibition of the OFC→CDS pathway significantly impaired contextual regulation of reward prediction (P < 0.001; stratified bootstrap test, see Methods for details). No significant effects of LED stimulation on DI were observed in the OFC→MDT or mCherry control groups (**Fig. 1F, Table S1**). Between-subject comparisons further revealed that the effect of LED stimulation in the OFC→CDS group differed significantly from both the OFC→MDT and control groups (Ps < 0.001), while no difference was observed between the OFC→MDT and control groups (P = 0.285; bootstrap test) (**Fig. S4**).

### OFC→cDS and OFC→MDT pathway inhibition produce asymmetric effects on the different target cues

In previous analyses, the four trial types were grouped in two broad categories: rewarded (A:X+, Y+) and nonrewarded (X-, A:Y-). Here we examined the effect of LED stimulation on each trial type individually (**Figs. 2 and S5**). For this, we fit LMMs separately for each group, including LED condition, Context, and Cue as within-subject factors —allowing the model to distinguish between all four trial types. For OFC→CDS pathway inhibition, this trial-level approach revealed that the impact of LED stimulation was dependent on specific cue-context combinations, i.e. trial identity (LED x Cue x Context: F(1, 5742.1) = 244.3626; P < 0.001). Post-hoc comparisons revealed that while LED stimulation consistently increased responding to nonrewarded trials (X− and A:Y−) and decreased responding to rewarded trials (A:X+ and Y+), the magnitude of these effects varied substantially across cues. Specifically, OFC→CDS pathway inhibition predominantly disrupted the contextual regulation of cue Y; LED stimulation significantly increased responding to the A:Y− trials (T(5742) = 17.361, p < 0.001, d = 0.9907) and significantly decreased responding to the Y+ trials (T(5742) = 9.830, p < 0.001, d = 0.7931). In contrast, the effects on cue X were more modest: LED stimulation significantly reduced responding to A:X+ trials (T(5742) = 3.768, p < 0.001, d = 0.304), while the increase in responding to X− trials did not reach significance (T(5742) = 1.627, p = 0.0885) (**Fig. 2A-C**). This asymmetric effect of OFC→CDS pathway inhibition across target cues was further supported by analyses of the Discrimination Index. Whereas previous analyses computed a single DI across trial types, here we calculated separate indices—DIX and DIY—to quantify the contextual modulation of cues X and Y, respectively. Inhibition of the OFC→CDS pathway led to a substantial reduction in DIY, indicating a strong disruption of contextual control over cue Y, whereas DIX showed only a modest decrease (**Figs. 2D and S5**). Together, these results indicate that OFC→CDS output is particularly critical for contextual regulation of cue Y —i.e. for resolving the A:Y- / Y+ discrimination.

**Fig. 2.**
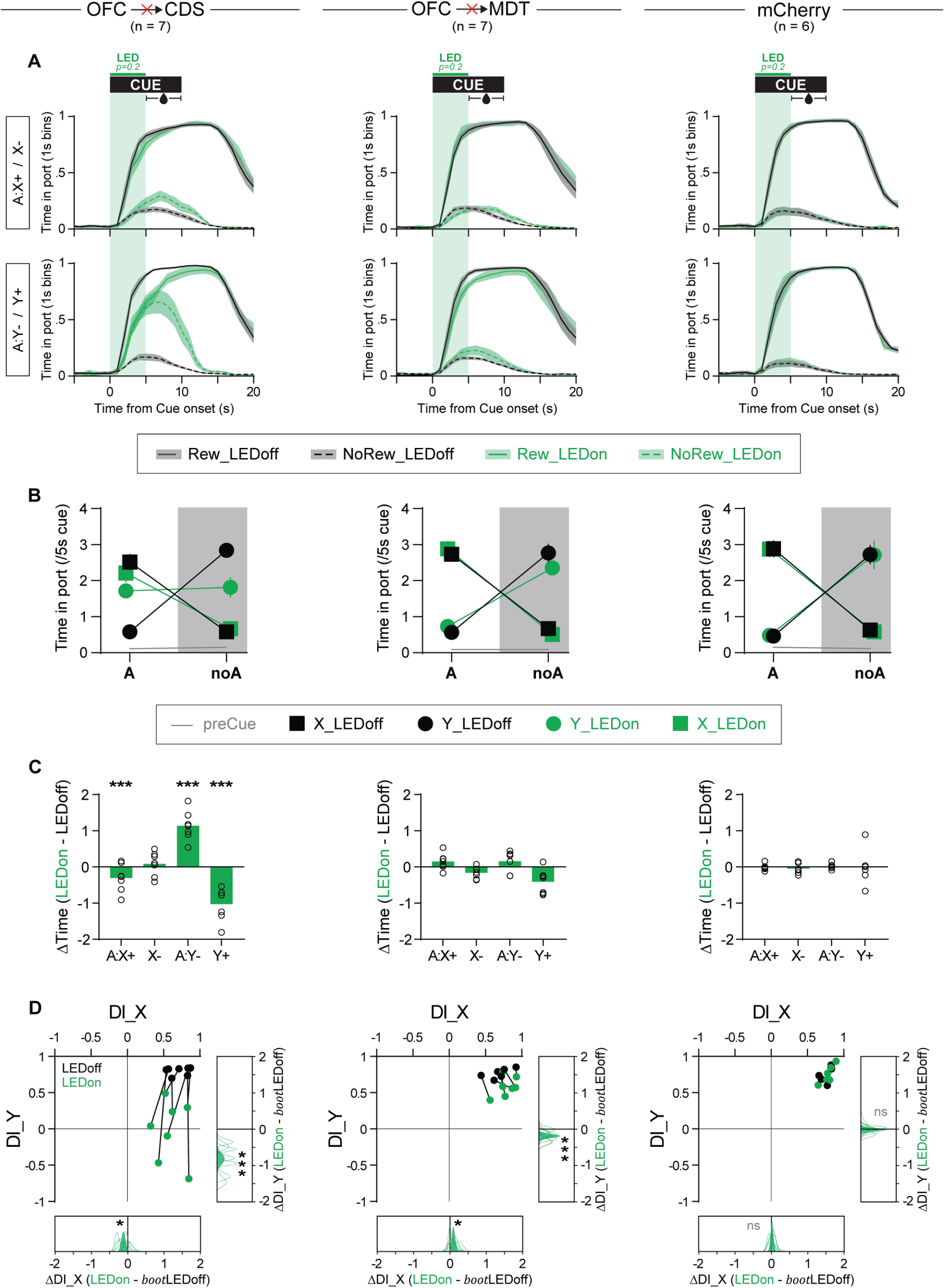
OFC→CDS and OFC→MDT pathway inhibition produce asymmetric effects across target cues. **A**. Port occupancy throughout cue presentation. Top: target cue X (i.e. context-gated discrimination A:X+ / X-). Bottom: target cue Y (i.e. context-gated discrimination A:Y-/ Y+). **B**. Mean port occupancy during the first 5 s of cue presentation (prior to reward delivery) for each trial type (A:X, X, A:Y, Y). **C**. Effect of LED stimulation on port occupancy expressed as a difference score (LEDon − LEDoff). Silencing OFC→CDS pathway increased responding to non-rewarded cues and decreased responding to rewarded cues. These changes were observed for both target cues but were substantially larger for cue Y (*** P < 0.001, LEDoff vs. LEDon, LMM post-hoc). In contrast, OFC→MDT silencing produced modest generalized effects, characterized by increased cue-evoked responding in Ctx. A and decreased cue-evoked responding in Ctx. noA, across all target cues (LED x Context interaction: P<0.001; LMM). **D**. auROC-based discrimination indices for target cue X (x-axis) and target cue Y (y-axis). Insets: effect of LED stimulation on discrimination performance expressed as ΔDI (LEDon − bootstrapped LEDoff). Distributions show bootstrapped ΔDI estimates for individual subjects (green lines) and the group-level estimate (green area). OFC→cDS pathway silencing disproportionately disrupted contextual regulation of responding to cue Y, effectively abolishing A:Y-/ Y+ discrimination, while producing only a modest impairment in contextual regulation of responses to cue X (A:X□ / X□). In contrast, OFC→MDT pathway silencing produced opposite effects across target cues, mildly disrupting contextual regulation of responses to cue Y while seemingly enhancing contextual regulation of responses to cue X (*: P<0.05; ***: P < 0.001, LEDoff vs. LEDon, bootstrap test). In m-Cherry (control) rats, cue-evoked responding and discrimination performance were unaffected by LED stimulation.

For OFC→MDT pathway inhibition —which appeared to have no overall effect when trial types were collapsed— analysis at the level of individual trial types revealed small but significant context-dependent effects. Although inhibition of this pathway did not impair context-dependent discrimination per se (LED × Cue × Context: F(1, 5746) = 2.32, p = 0.128), it did produce a more global, context-dependent shift in cue-evoked responding (LED x Context: (F(1, 5746) = 26.32, p < 0.001). Specifically, inhibition of this pathway caused a generalized increase of cue-evoked responding in the presence of context A (A:X+ and A:Y-collapsed, T (5746) = 2.413; P = 0.0159; d = 0.119) and a generalized decrease of cue-evoked responding in the absence of context A (X- and Y+ collapsed, T (5746) = 4.843; P < 0.001; d = 0.239) (**Fig. 2A-C**). As a result, the discrimination index for cue Y was slightly decreased (DIY_LEDoff vs DIY_LEDon: P < 0.001) following OFC→MDT pathway inhibition, whereas the discrimination index for cue X was slightly increased (DIX_LEDoff vs DIX_LEDon: P = 0.02036) (**Fig. 2D**). These results suggest that OFC→MDT inhibition does not disrupt contextual gating process but instead reveals subtle latent imbalance in the relative value of the two contexts.

In mCherry control rats, LED stimulation had no effect on cue-evoked responding (no main effect or interaction with LED stimulation: (F(1, 6393) < 1.6002; Ps > 0.2059). Moreover discrimination indexes remained unchanged following LED stimulation (**Fig. 2**).

### Phasic inhibition of OFC→cDS pathway has little effect on simple (linear) discrimination tasks

Our results thus far show that inhibition of the OFC→CDS pathway profoundly disrupts the regulation of reward seeking in a context-dependent discrimination task —a form of nonlinear discrimination in which reward predictions depend on a nonlinear combination of contextual and target cues. To assess the specificity of this effect, we next examined the consequences of this manipulation in rats trained on simpler, linear discrimination tasks (**Figs. 3 and S5**).

**Fig. 3.**
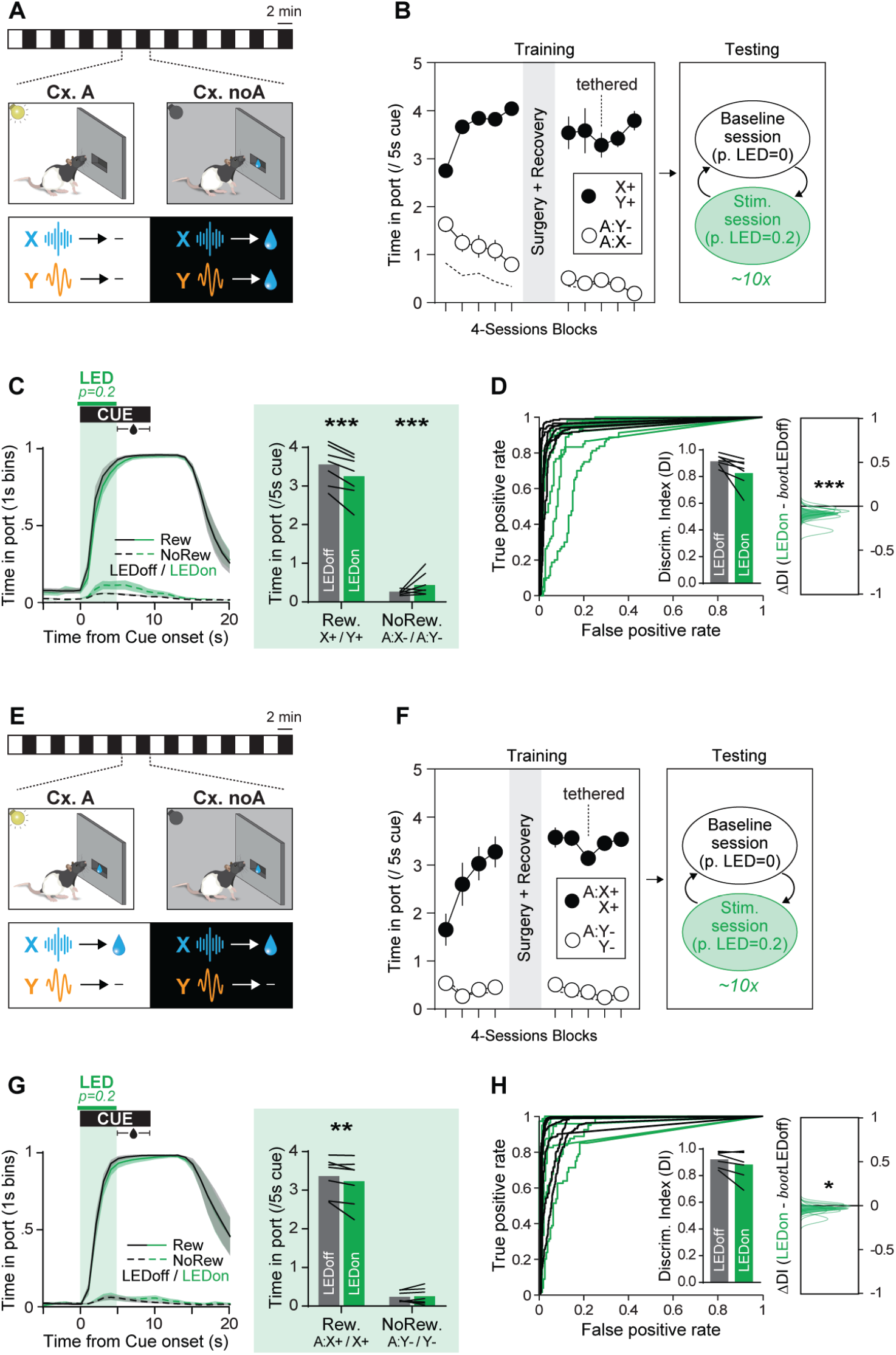
Phasic inhibition of OFC→CDS pathway has little effect on linear discrimination tasks. **A-D**. Effects of OFC→CDS pathway silencing in strictly negative occasion setting task. **A**. Negative occasion setting task: auditory target cues X and Y were followed by reward when presented alone, but not when presented in compound with the visual contextual cue A (A:X-/ X+ / A:Y-/ Y+). **B**. Experimental timeline. Following initial training, rats underwent surgery and post-surgical retraining. During subsequent stimulation sessions, LED stimulation was applied on 20% of trials. Stimulation sessions alternated with baseline sessions during which no stimulation was delivered. **C**. Port occupancy throughout cue presentation. Histograms summarize mean port occupancy during the first 5 s of the cue, before reward delivery. Lines represent individual subjects (*** P < 0.001, LEDoff vs. LEDon, LMM post-hoc). **D**. ROC curves and auROC-based discrimination index. Insets: effect of LED stimulation on discrimination performance expressed as ΔDI (LEDon − bootstrapped LEDoff). Distributions show bootstrapped ΔDI estimates for individual subjects (green lines) and the group-level estimate (green area). OFC→cDS pathway silencing produced small but significant reduction in discrimination performance (*** P < 0.001, LEDoff vs. LEDon, bootstrap test). **E-H**. Effects of OFC→CDS pathway silencing in simple auditory discrimination task. **E**. Simple auditory discrimination task. Only target cue X is rewarded, independent of background visual context (A:X+ / X+ / A:Y-/ Y-). **F**. Experimental timeline. **G**. Port occupancy throughout cue presentation. Histograms summarize mean port occupancy during the first 5 s of the cue, before reward delivery. Lines represent individual subjects (** P < 0.01, LEDoff vs. LEDon, LMM post-hoc). **H**. ROC curves and auROC-based discrimination index Insets: effect of LED stimulation on discrimination performance expressed as ΔDI (LEDon − bootstrapped LEDoff). Distributions show bootstrapped ΔDI estimates for individual subjects (green lines) and the group-level estimate (green area). OFC→cDS pathway silencing produced small but significant reduction in discrimination performance (* P < 0.05, LEDoff vs. LEDon, bootstrap test).

First, in light of the dramatic effects of OFC→CDS silencing on the contextual regulation to cue Y —the cue subject to negative contextual gating (A:Y- / Y+)— we examined the impact of this manipulation in rats trained on a strictly negative occasion setting task, in which the contextual cue A always signaled the absence of reinforcement (A:X- / X+ / A:Y- / Y+) (**Fig. 3A-D**). In this task, both target cues were functionally equivalent and thus collapsed for analysis. LMM revealed that inhibition of the OFC→CDS pathway disrupted negative occasion setting (Context × LED interaction: F(1, 7495) = 77.5431; P < 0.001) by increasing responding to nonrewarded cues (T(7495) = 6.434; P < 0.001) and decreasing responding to rewarded cues (T(7495) = 6.236; P < 0.001). This impairment was also reflected in a significant reduction of the Discrimination Index following OFC→CDS pathway silencing (DI_LEDon vs. DI_LEDoff: P < 0.001) (**Fig. 3 C-D**).

Because each context cycle contained two consecutive trials, in this strictly negative occasion-setting task, the rewarded and nonrewarded trials necessarily came in pairs (a 1st rewarded trial was always followed by 2nd rewarded trial, and likewise a 1st nonrewarded trial was always followed by 2nd nonrewarded trial). Reward delivery tends to transiently elevate responding on the subsequent trial —a “momentum” effect previously characterized in this task structure^16^. Therefore, in this strictly negative occasion-setting task, such outcome-driven momentum could incidentally enhance discrimination performance on the second trial of each cycle. Whereas responses to the first cue must rely solely on contextual information, responses to the second cue may additionally benefit from residual outcome-driven momentum, potentially masking the effects of OFC→CDS silencing on those trials. To account for this potential confound, we fit an additional LMM including trial number within the context cycle as a factor. This analysis revealed that the effect of OFC→CDS pathway silencing depended jointly on the context and the trial number in this context (LED x Context x Trial number: F(1, 4561.1) = 8.94, p = 0.0028). Specifically, LED stimulation disrupted contextual regulation on first cues by increasing responding to nonrewarded cues and reducing responding to rewarded cues (Ts (4561) > 4.443; Ps<0.001), but it had no significant effect on the second cues in a context cycle (Ts(4561)<1.871; P>0.0614). This suggests that the OFC→CDS pathway contributes to behavioral regulation when responding depends on context-sensitive predictions rather than recent outcome history^34^.

Even when accounting for the sequence effect by restricting analyses to first cues, the impact of OFC→CDS inhibition on contextual regulation in the strictly negative occasion-setting task remained modest (effect size: ds < 0.387). This contrasted sharply with the profound disruption caused by the same manipulation to negative contextual gating (A:Y−/Y+) in rats trained on the nonlinear context-dependent discrimination task (ds > 0.793). The difference in disruption magnitude was confirmed statistically by comparing the stimulation-induced change in discrimination index (ΔDI) between groups (nonlinear ΔDI_Y vs. linear ΔDI: p < 0.001).

These results indicate that what appears to be the same contextual discrimination problem (A:cue- / cue+) does in fact engage partially distinct neural substrates when embedded in a linear versus a nonlinear discrimination task.

Finally, we examined the effect of OFC→CDS inactivation in a simple auditory discrimination task, where cue X was always rewarded, cue Y never rewarded, and the background context was irrelevant (A:X+ / X+ / A:Y− / Y−) (**Fig. 3E-H**). Inhibition of the OFC→CDS pathway caused a small but significant disruption in discrimination performance (Cue x LED interaction: F(1, 8150) = 8.6764; P = 0.003233) driven by a modest reduction of responding to rewarded cues ((T(8150) = 3.178; P = 0.0015) causing modest decrease in the discrimination index (P= 0.0357) (**Fig. 3G-H**). The magnitude of this effect was very small (d= 0.152), once again contrasting with the profound disruption caused by the same manipulation in rats trained in the nonlinear context-dependent discrimination task (**Fig. S5**).

Together, these results indicate that OFC→CDS activity plays only a minor role in regulating reward seeking during simple, linear discriminations, but becomes critical when contextual gating is required.

## DISCUSSION

The orbitofrontal cortex (OFC) is essential for the contextual regulation of reward prediction and reward seeking^17–20^. Yet, the circuit-level mechanisms through which the OFC exerts this context-dependent control have remained elusive. Here, we examined the contribution of two major OFC efferent pathways — projections to the central dorsal striatum (OFC→CDS) and to the mediodorsal thalamus (OFC→MDT)— in the contextual gating of cued reward predictions. Using temporally precise optogenetic inhibition of OFC terminals in rats performing a context-dependent discrimination task, we identified dissociable, cue- and pathway-specific roles for these OFC outputs.

Silencing the OFC→CDS pathway profoundly disrupted the contextual regulation of reward seeking: rats increased responding to nonrewarded cue/context configurations and decreased responding to rewarded configurations. Although both target cues were affected, the disruption was especially pronounced for the negatively gated component (A:Y− / Y+). In contrast, inhibition of the OFC→MDT pathway produced more modest and qualitatively different effects. Indeed, silencing of this pathway did not abolish contextual gating per se; instead, it caused a modest global increase in cue-evoked responding in context A and a decrease in context noA, regardless of cue identity. This behavioral pattern suggests that OFC→MDT activity normally constrains latent imbalances in contextual value; when silenced, pre-existing differences between contexts are amplified.

Importantly, the contribution of OFC→CDS was relatively specific to hierarchical, context-gated predictions (i.e. predictions that engage nonlinear cue/ context configurations). Indeed, silencing OFC→CDS pathway had modest to minimal impact in simpler linear discrimination tasks such as strictly negative occasion setting, or simple auditory cue discrimination. Thus, OFC→CDS appears particularly critical when task performance depends on hierarchical associative structure, rather than when behavior can be supported by simple binary associations.

A striking feature of these results is the asymmetric impact of OFC pathway inhibition across the two discrimination components: positive gating (A:X+ / X-) and negative gating (A:Y-/ Y+). Silencing the OFC→CDS pathway virtually abolished the negative gating component of the task, while producing only a modest disruption on the positive gating component. Silencing the OFC→MDT pathway slightly impaired negative contextual gating but, surprisingly, improved positive gating —a notable effect given that baseline discrimination performance was already high and near ceiling. This cue-asymmetry provides a useful window to further delineate the distinct contributions of the OFC→CDS and OFC→MDT pathways to the contextual regulation of behavior. Although previous studies have implicated the OFC in modulating sensory processing, including auditory perception^35–40^, the present cue-asymmetry indicates that rats retained the ability to discriminate both cues and contexts despite OFC→CDS or OFC→MDT pathway silencing. Together with the modest effects of OFC→CDS inhibition in the negative occasion-setting and simple discrimination tasks, this pattern demonstrates that the disruption observed following OFC pathway silencing cannot be explained by sensory deficits or generalized motivational impairments.

To better understand the possible origin of these cue-asymmetric effects, we implemented a connectionist model of associative learning —building on architectures developed to explain occasion setting and other nonlinear discriminations^41,42^ (**Fig. 4**). Briefly, the model consists of a simple feedforward network in which incoming stimuli connect to a reward-prediction output through a layer of hidden units. These include elemental units —each receiving input from a single stimulus— and configural units receiving convergent input from all stimuli. In this architecture, context can influence reward prediction in two distinct ways: through linear, additive contributions (context acting a regular cue via context-outcome associations) and through nonlinear configural representations (context acting as a gatekeeper of cue-evoked associative memories)^5,6^. Critically, configural units are assigned lower baseline activity, making them harder to recruit (**Fig. 4A**). As a result, the network (trained via backpropagation^43^) preferentially adopts elemental, additive solutions early in learning, with configural units engaged only when elemental strategies fail to achieve accurate predictions after extended training. Under these assumptions (segregated elemental/configural pathways and early dominance of the elemental path) we found that the network solved the context-dependent discrimination using a heterogeneous mixture of associative strategies. Specifically, the negative gating component (A:Y− / Y+) relied heavily on the configural pathway, whereas the positive gating component (A:X+ / X−) was solved primarily through elemental associations with only moderate configural involvement (**Fig. 4D**). Synthetic “lesions” (or silencing) of the configural pathway therefore produced dramatic impairment in the negative contextual gating and a more modest disruption of positive gating —closely mirroring the behavioral effects of OFC→CDS silencing (**Fig. 4E**). The model also recapitulated task specificity: configural pathway disruption produced maximal impairment in the nonlinear discrimination, moderate effects in negative occasion setting, and little to no effect in simple discrimination (**Fig. S6**). Collectively, these findings support the interpretation that the OFC→CDS pathway conveys, or enables, configural cue/context representations required for context-gated associative predictions.

**Fig. 4.**
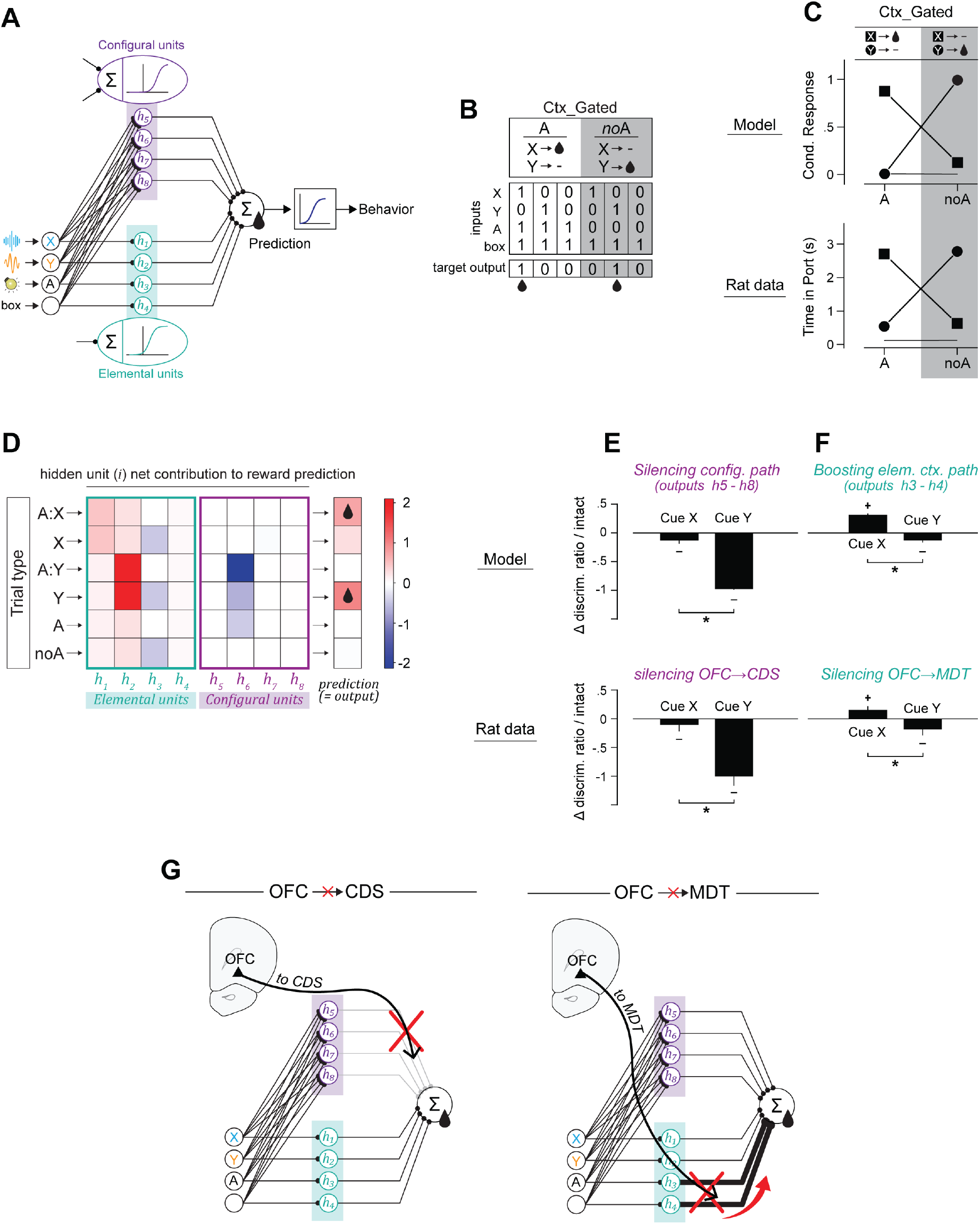
Connectionist model accounts for pathway-specific OFC contributions. **A**. Model architecture. The model consists of a feedforward network in which stimulus inputs connect to a reward-prediction output through hidden units. Hidden units include elemental units, each receiving input from a single stimulus, and configural units, which receive convergent input from all stimuli. Configural units have lower resting activity, biasing learning toward the elemental pathway, with the configural pathway recruited when elemental strategies fail to achieve accurate predictions. **B**. Context-dependent behavioral task formalized as inputs-target outputs matrix for training via backpropagation **C**. Network performance (top) and rat behavior (bottom) after training to criterion. **D**. Hidden-unit contributions to outcome prediction. The contribution of each hidden unit (i) was computed as A_i_ × W_i_→out (A_i_: unit activity; W_i_→out: weight to output). Configural units contribute predominantly to the A:Y-/ Y+ component, with little involvement in A:X+ / X-. **E**. Simulated disruption of configural pathway. Reducing the influence of the configural pathway strongly disrupts network discrimination for target cue Y (A:Y-/ Y+) and produces more modest effects for target cue X (A:X+ / X-) (top), closely mirroring the effects of OFC→CDS silencing observed in rats (bottom). **F**. Simulated enhancement of elemental pathway. Amplifying contextual elemental inputs magnifies small context-value imbalances, moderately disrupting discrimination for target cue Y (A:Y-/ Y+) while improving contextual regulation for target cue X (A:X-/ X+) (top), closely mirroring the effects of OFC→MDT silencing observed in rats (bottom). **G**. Proposed contributions of OFC output pathways. OFC→CDS enables context-gated predictions by conveying configural representations. OFC→MDT limits the influence of direct context–outcome associations in reward prediction. Discrimination performance was quantified using the discrimination ratio: (Rrewarded - Rnonrewarded) / (Rrewarded + Rnonrewarded). +/−: P < 0.05 significant change from zero (one-sample t-test). *: P < 0.05 (paired t-test). Model simulation: n = 20; Rat data: n = 7.

How to interpret the effect of OFC→MDT silencing? Within the same connectionist framework, the effects of OFC→MDT silencing were reproduced by increasing the gain of contextual elemental units —those dedicated to representing the houselight (Ctx. A) and the broader conditioning chamber (“box”). Amplifying these units effectively increased the additive influence of elemental contextual value, thereby magnifying subtle imbalances between contexts. This manipulation recapitulated the behavioral pattern observed following OFC→MDT silencing: generalized increase of responding in context A and generalized decrease of responding in context noA, resulting in improved A:X+ / X− discrimination and reduced A:Y− / Y+ discrimination (**Fig. 4F**). These results suggest that contextual elemental representations are normally constrained —potentially via OFC→MDT signaling— a regulatory mechanism not explicitly implemented in the current model.

The OFC–striatal pathway has been implicated in compulsive and inflexible behaviors, yet its precise contribution remains debated. Both hyperactivation and hypoactivation of the OFC→CDS pathway have been associated with maladaptive responding in humans and rodents^26,33,44–46^. Our findings help reconcile these observations by showing that OFC→CDS activity does not exert a simple inhibitory or excitatory influence over behavior, but instead regulates behavior according to context-gated (hierarchical) associative structures. Within this framework, hyperactivity in the pathway could promote compulsive behavior by enforcing rigid contextual representations (as proposed in obsessive– compulsive disorder) whereas hypoactivity could allow behavior to default to context-invariant, stimulus-driven responding (as proposed in addiction). Maladaptive, compulsive behavior may therefore arise from dysregulated contextual gating mechanisms^3^.

Several important questions remain. First, what specific information is encoded by OFC neurons projecting to CDS and MDT? Do OFC→CDS neurons explicitly encode configural, context-dependent predictions, or do they instead modulate the influence of such representations computed elsewhere? Likewise, how do OFC→MDT neurons normally constrain the impact of elemental contextual value on behavior? Projection-specific recording during context-gated discrimination will be necessary to address these questions. Second, how does OFC input shape striatal activity to implement contextual control? One possibility is that OFC→CDS projections provide top-down modulation of cue-evoked activity in the striatum, dynamically biasing the relative engagement of direct- and indirect-pathway medium spiny neurons. Such modulation could occur via direct synapses onto striatal medium spiny neurons, or indirectly through OFC projections to striatal cholinergic interneurons, which exert powerful control over cue-evoked activity and motivated behavior^47–49^. Dissecting these mechanisms will require simultaneous monitoring/manipulation of OFC terminals and identified striatal cell populations.

Context-dependent predictions are fundamental to adaptive, flexible behavior, allowing animals to extract context-appropriate meaning from ambiguous cues and react in a context-appropriate manner. Our findings indicate that the OFC orchestrates such flexibility through at least two dissociable efferent pathways with distinct functional roles. The OFC→CDS projection enables hierarchical, context-sensitive predictions to guide behavior. In contrast, the OFC→MDT pathway makes a more modest yet complementary contribution by limiting the influence of direct context–outcome associations, thereby promoting hierarchical gating over direct context-driven responding. Together, these results reveal a division of labor across OFC efferent pathways that enables adaptive, context-appropriate behavior.

## METHODS

### Subjects

Male and female Long-Evans rats (Charles River), aged 8-10 weeks upon arrival, were housed in a temperature-controlled vivarium with a 12-hr light/ dark cycle, initially in same-sex pairs and later singly housed after surgery. Rats were mildly food restricted to maintain ∼95% of age-matched free-feeding weight, except during the pre- and post-surgery recovery period when food was available ad libitum. Water was always available in the home cage. All experiments were conducted during the light phase. Because acquisition of the context-dependent discrimination task required substantially longer training, rats assigned to this condition began training at a younger age (immediately upon arrival), whereas rats trained on other tasks began later (∼4 months old). This ensured that all rats were of comparable ages during the critical phases of the experiment. All experimental procedures were conducted in accordance with UCSB Institutional Animal Care and Use Committees and the US National Institute of Health guidelines.

### Apparatus

Behavioral training was conducted in 12 identical conditioning chambers, each enclosed in a sound-attenuating cubicle (Med Associates). A fan mounted on the cubicle provided ventilation and low-level background noise. Each chamber was equipped with two auditory cue devices: a 2.5Hz clicker mounted on the front panel, and a white noise generator on the back panel (∼76dB each). Two ceiling-facing lights, positioned on the front and back walls, provided diffuse illumination and were used to manipulate the visual context. Sucrose solution (15% w/v) was delivered in a recessed reward port, located in center of the front panel, via a syringe pump located outside the sound-attenuating cubicle. Head entry into the reward port was detected by interruption of an infrared beam. Training sessions included rats of both sexes, tested concurrently in sex-specific chambers.

### Discrimination training

Behavioral tasks were adapted from Peterson et al.^16,17^. Rats were then trained daily (5-6 days/week) in 2-h sessions during which the visual context alternated every 2 min between context A (flashing house lights, 0.25 s on/0.25 s off) and context noA (all lights off). Within each context, two 10-s auditory cues, X or Y (white noise or clicker, counterbalanced), were presented individually in pseudorandom order. This generated four trial types: A:X / X / A:Y / Y (120 trials in 2h session; ITI = 50±20s).

#### Context-dependent discrimination

the predictive status of each cue was informed by the context. Specifically, cue X was rewarded with sucrose only in the presence of the context A, and cue Y was rewarded only in the absence of context A (A:X+ /X- /A:Y- / Y+). For the first 30 training sessions all trials were in equal proportion. Then, to discourage indiscriminate responding and promote discrimination learning, the proportion of rewarded trials was reduced to account for one third of all trials, evenly distributed across contexts (20 A:X+ / 40 X- / 40 A:Y- / 20 Y+).

#### Negative occasion-setting

both cues were rewarded only in absence of context A (A:X- / X+ / A:Y- / Y+). For the first 12 training sessions all trials were in equal proportion. Then, the proportion of rewarded trials was reduced to match the context-dependent discrimination task (40 A:X- / 20 X+ / 40 A:Y- / 20 Y+). Note that this adjustment required uneven exposure to the two contexts, resulting in context A occasionally extending for more than 2 min.

#### Simple auditory discrimination

cue X was always rewarded and cue Y was never rewarded, regardless of context (A:X+ / X+ / A:Y-/ Y-). For the first 12 training sessions all trials were in equal proportion. Then, the proportion of rewarded trials was reduced to match the context-dependent discrimination task (20 A:X+ / 20 X+ / 40 A:Y-/ 40 Y-).

All rewards consisted of 0.2mL sucrose (15% v/w) delivered over 5s, starting 5s after cue onset. Training initially followed strictly Pavlovian contingencies, with reward delivery independent of behavior. After surgery and once stable discrimination was reestablished, reward delivery required the rat’s presence in the port during the second half of the cue. This prevented potential leftover sucrose from contaminating performance, which is critical for optogenetic experiments where neural manipulations might have increased response omission rates.

### Surgeries

Rats were anesthetized with isoflurane (5% induction, 1–2.5% maintenance), placed in a stereotaxic frame, and maintained at ∼37 °C. Bilateral OFC injections (0.8 µL/site) of AAV8-hSyn-eNpHR3.0-mCherry (GVVC-AAV-147) or AAV8-hSyn-mCherry (Addgene 114472-AAV8) were made at the following coordinates: anterior–posterior (AP) +4.0 mm from bregma, medial– lateral (ML) ±2.2 mm from midline, and dorsal–ventral (DV) −5.0/−4.5 mm from skull surface. Virus was delivered through 33-gauge injectors at a rate of 0.1 µL/min; injectors remained in place for 5 min post-infusion to permit diffusion. In the same surgery, rats were implanted bilaterally with optic fibers (300 µm core; RWD R-FOC-BF300C-39NA) targeting the CDS (AP +0.8 mm, ML ±2.7 mm, DV −4.6 mm) or MDT (AP −2.8 mm, ML ±1.75 mm, DV −5.4– mm at a 10° angle). Implants were secured with skull screws and black dental acrylic (Lang Dental 1530BLK). Rats received carprofen (5 mg/kg, p.o.) once daily for 4 days starting 2 h before surgery and were allowed to recover for at least 1 week before resuming training. Optogenetic inhibition sessions were conducted >6 weeks post-surgery to ensure sufficient viral expression at the targeted regions.

### Optogenetic Inhibition

Rats were habituated to the tethering system for at least 12 days (extended if required to re-establish pre-surgery discrimination performance). The tethering system consisted of a lab-built bifurcating Y-shaped optical patch cord (0.63 NA, 500 µm core diameter, encased in a flexible stainless steel jacket) suspended to a balancing arm and connected to a LED light source via a fiber-optic rotary joint. During optogenetic inactivation sessions, the LED (540 nm, 8–10 mW measured at the fiber tip) was activated for 5 s, beginning 200 ms before cue onset, overlapping the reward anticipation period. Each session included 120 trials (40 rewarded, 80 non-rewarded), with LED delivery on 20% of trials (8 rewarded, 16 non-rewarded), equally distributed across contexts. Each rat underwent ∼10 stimulation sessions (∼1200 total trials, ∼240 LED-on trials), which alternated with regular behavioral sessions without LED delivery.

Because of the importance of visual stimuli in this task, several measures were taken to prevent LED light leakage: (i) fibers were encased in opaque metal ferrules, (ii) the length of exposed fiber between skull and ferrule was minimized (<1 mm), (iii) implants were secured to skull with high-opacity black dental cement, and (iv) patch cords were connected to implants using opaque metal sleeves covered with heat-shrink tubing. Under these conditions, LED light delivery was not visually detectable to human observers.

### Histological verification

Rats were deeply anesthetized with Euthasol and transcardially perfused with PBS followed by 4% paraformaldehyde. Brains were extracted, postfixed in 4% paraformaldehyde for 24h, cryoprotected in 30% sucrose for >3 days, and sectioned at 40µm on a cryostat. Coronal sections were mounted on glass slides and coverslipped with a DAPI-containing mounting medium (Vectashield-DAPI, Vector Laboratories). Images were acquired using a fluorescence microscope (Keyence BZ-X800). Viral (mCherry) expression in the OFC was verified in all rats; mCherry expression and fiber tip placements were also verified in the CDS or MDT depending on experimental group.

### Statistical Analysis

#### Time in Port

Cue-evoked reward seeking was quantified as the time spent in the reward port during the first 5 s of cue presentation, prior to reward delivery. Time in port was analyzed with linear mixed-effects models (LMMs; lme4 package in R) using restricted maximum likelihood (REML) estimation and Satterthwaite’s approximation for degrees of freedom. Group was included as a between-subject factor, and Trial (or Cue × Context) and LED stimulation were included as within-subject factors, with Subject as a random effect. Post hoc pairwise comparisons of estimated marginal means (emmeans package in R) were conducted, and effect sizes (Cohen’s d) were calculated for significant contrasts.

#### Discrimination Index

To facilitate group comparisons, we calculated a discrimination index (DI) for each animal. The DI was based on the area under the receiver operating characteristic curve (auROC) defined by the distributions of responding to rewarded versus non-rewarded cues (perfcurve function in Matlab). For interpretability, auROC values were rescaled from −1 to 1 using the formula DI = 2 × (auROC − 0.5), where 1 indicates perfect discrimination (responses to rewarded cues always exceeding responses to non-rewarded cues), 0 indicates no discrimination, and −1 indicates inverse discrimination. This empirical auROC-based DI makes no distributional assumption and is particularly adapted to this behavioral task in which responses are not normality distributed50,51.DIs were computed under both LED_on and LED_off conditions, and the difference between these conditions was used to quantify the effect of optogenetic manipulation on discrimination performance.

To assess the effect of LED stimulation on discrimination performance, we used a stratified bootstrapping approach. For each rat, 50,000 resampled datasets were generated from LEDoff trials, with each resample matched in size to the LEDon condition (i.e., containing the same number of rewarded and non-rewarded trials). A DI was computed for each resample (using the auROC approach described above), and the difference between the observed DI_LEDon and the resampled DI_LEDoff was calculated:

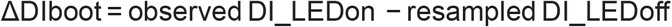

This procedure produced a bootstrap distribution of 50,000 ΔDI values per subject, providing an estimation of the effect of LED stimulation on DI and the likelihood that any observed deviation could be explained by subsampling of LEDoff trials (since only 20% of all trials were LEDon). To obtain group-level estimates, we resampled across subjects by drawing (with replacement) one ΔDIboot value from each subject’s distribution and averaging across all subjects; this procedure was repeated 50,000 times to generate a group-level bootstrap distribution of ΔDI.

To assess the effect of LED stimulation on discrimination performance within each group, we tested the null hypothesis that LED stimulation had no effect on DI (ΔDI = 0). Two-tailed p-values were defined by the proportion of bootstrap ΔDI values that crossed or exceeded zero, representing the probability that the observed difference between LEDon and LEDoff DI could be explained by chance resampling of LEDoff trials alone.

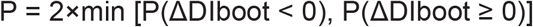

Group differences in the effect of LED stimulation (ΔDI) were assessed using two complementary approaches. First, we performed Welch’s ANOVA on the observed subject-level ΔDI values, followed by post hoc pairwise Welch’s t-tests with p-values adjusted using the Holm– Šidák procedure to control for multiple comparisons. Second, we extended the bootstrap framework to between-group contrasts by leveraging the group-level bootstrap distributions of ΔDI. At each iteration, one ΔDI value was randomly drawn from each group’s distribution, and pairwise differences were computed (ΔΔDI = ΔDI_group1 − ΔDI_group2). Repeating this 50,000 times generated empirical sampling distributions of ΔΔDI for every pairwise comparison. From these distributions, we derived two-tailed p-values where p reflects the probability that the observed group difference could have arisen from random variation in trial- and subject- sampling. Because this procedure propagates trial- and subject-level uncertainty, no additional corrections for multiple comparisons was applied. Both approaches produced convergent results (Table S2). In the main text, we report the bootstrap p-values. All tests were two-tailed, significance was assessed against a type I error rate of 0.05. Statistical analyses were conducted with Matlab (R2024b) and R (version 4.5.1).

Criterion for inclusion in the experiment was arbitrarily defined as responding in rewarded trials at least 1.75 greater than responding in nonrewarded trials over a 4-session period, in baseline conditions (prior to optogenetic manipulations). Two rats trained in the context-dependent discrimination task were excluded for failing to reach this this criterion within 90 sessions. Six more rats were excluded due to insufficient/ incorrect viral expression or misplaced fiber implants. LED light delivery in mCherry-expressing rats had no detectable effect in either CDS or MDT. To minimize animal use, and given the extensive training required for context-dependent discrimination, these rats were pooled into a single mCherry control group for statistical analyses. Although both male and female rats were included, group sizes per sex were small (n = 3–4 M or F per group), precluding meaningful statistical inference regarding sex differences. Accordingly, sex was not included as a factor in the final analyses, and results are reported collapsed across sex. Subject-level information, including sex, is provided in the accompanying data table (Table S1) and Figs. S3, S4, and S5.

### Connectionist model

The proposed model borrows largely from previous connectionist models of associative learning^41,42^ and consists of a simple feedforward neural network with three layers: (1) an input layer representing the target cues X and Y, the contextual cue A, and an additional unit capturing the background sensory stimuli continuously present in the conditioning chamber; (2) a hidden layer; and (3) a single output unit representing reward expectation. Net input to hidden units is transformed into a propagated signal (range: 0–1) via a logistic activation function. A negative bias term is subtracted from the net input to each hidden unit, ensuring that units these units remain relatively silent in absence of excitatory inputs.

Consistent with previous models, we make two key assumptions: Assumption #1. Hidden units are segregated into two classes: elemental units, which receive input from a single stimulus, and configural units, which receive convergent input from all stimulus. Assumption #2. Configural units have a larger negative bias than elemental units (−4 vs. −2.5), resulting in lower resting activity (0.02 vs. 0.08, respectively). Together, these assumptions bias the network toward elemental solutions early in training. Configural solutions are progressively recruited only if elemental strategies fail to accurately predict trial outcomes.

Behavioral tasks were formalized as matrices of input patterns and target outputs corresponding to the specific task contingencies. Following weights initialization with small random values (0–0.01), the network was trained using the standard backpropagation algorithm (learning rate = 0.08). All simulations were conducted in MATLAB R2024b.

To transform these network-generated predictions into ‘observable’ behavioral responses, we applied a sigmoidal transformation:

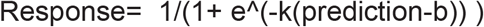

Where k determines the steepness of the sigmoid (here, k = 10) and b defines the inflection point (here, b = 0.5). This function has been shown to provide a parsimonious and principled transformation of associative predictions into observable behavior^52,53^. Training was terminated once the model satisfied the criterion that responding to each rewarded target cue was at least tenfold greater than the response to the corresponding nonrewarded cue (e.g., A:X+ ≥ 10* X- and Y+ ≥ 10* A:Y-in the context-gated task). This conservative criterion was chosen to approximate a near-asymptotic state of the network and aligns with the average discrimination levels observed in rats after excluding outlier trials (defined as responses > 2 SD from the mean) which correspond to an estimate of optimal, “noise-free”, behavioral performance.

#### Synthetic lesions within the connectionist model

Following training, we simulated pathway-specific silencing by selectively manipulating connection weights (W) within the network, to parallel experimental optogenetic manipulations.

To model disruption of configural representations —intended to approximate silencing of OFC→cDS pathway supporting context-gated (hierarchical) integration— we selectively attenuated the projections from configural hidden units to the output layer:

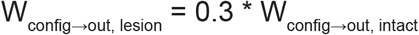

This manipulation reduces the influence of configural (stimulus × context) representations on reward prediction while preserving elemental stimulus processing.

To model an increase in the influence of elemental contextual representations —intended to approximate silencing of the OFC→MDT pathway and the resulting amplification of elemental context value— we selectively increased the contribution of elemental contextual units to the output layer:

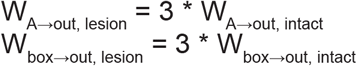

Because the network produces a single scalar response value per trial type, auROC-based discrimination indices are not applicable. Therefore, the impact of synthetic lesions on network performance was quantified using a discrimination ratio computed from the model responses to rewarded and nonrewarded trial types:

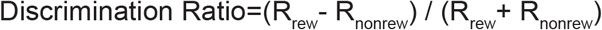

This discrimination ratio was computed for both the intact and lesioned networks, and the impact of the lesion was quantified as the proportional change in discrimination ratio. For direct comparison with empirical data, the same discrimination ratio was computed from rat behavior data. This parallel analysis confirms that the principal conclusions are robust and do not depend on the specific discrimination metric used (auROC vs. discrimination ratio).

## SUPPLEMENTAL MATERIAL

**Fig. S1:**
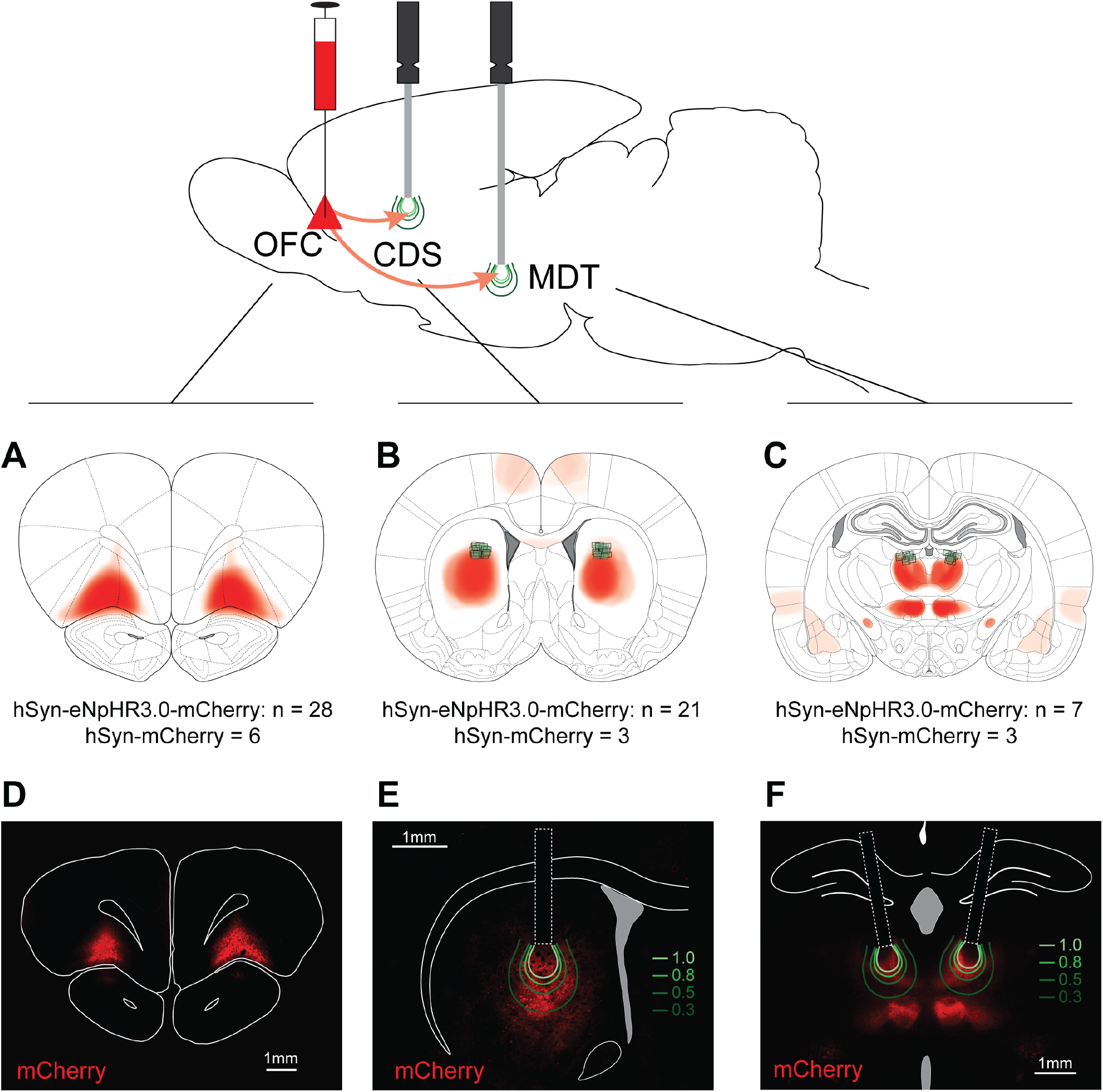
Virus expression and estimated LED light propagation. Viral vector for the expression of the inhibitory opsin eNpHR3.0 (hSyn-eNpHR3.0-mCherry) or the control transgene mCherry (hSyn-mCherry) was injected bilaterally in the orbitofrontal cortex (OFC). Optical fibers were implanted bilaterally, aimed at the central dorsal striatum (CDS), or the mediodorsal thalamus (MDT) to enable light-mediated activation of the opsin and inhibition of OFC projections to these target regions. **A-C**. Reconstructed representation of viral expression (combined across mCherry and hM4Di-mCherry subjects) in the OFC (A), and in the projections targets CDS (**B**) and MDT (**C**). In addition to those primary target regions, robust mCherry expression was also observed in the submedius thalamus. Green squares indicate the ventral tip of optical fiber placements in the CDS and MDT. **D-F**. Representative examples of virus expression in the OFC (**D**) and combined virus expression with fiber placement in the CDS (**E**) and MDT (**F**). Green contours depict estimated light iso-intensity lines corresponding to 100%, 80%, 50%, and 30% of maximum light intensity at the fiber tip. Light spread estimates are based on the direct visualization approach described in Johnson et al. (2021)^54^.

**Fig. S2:**
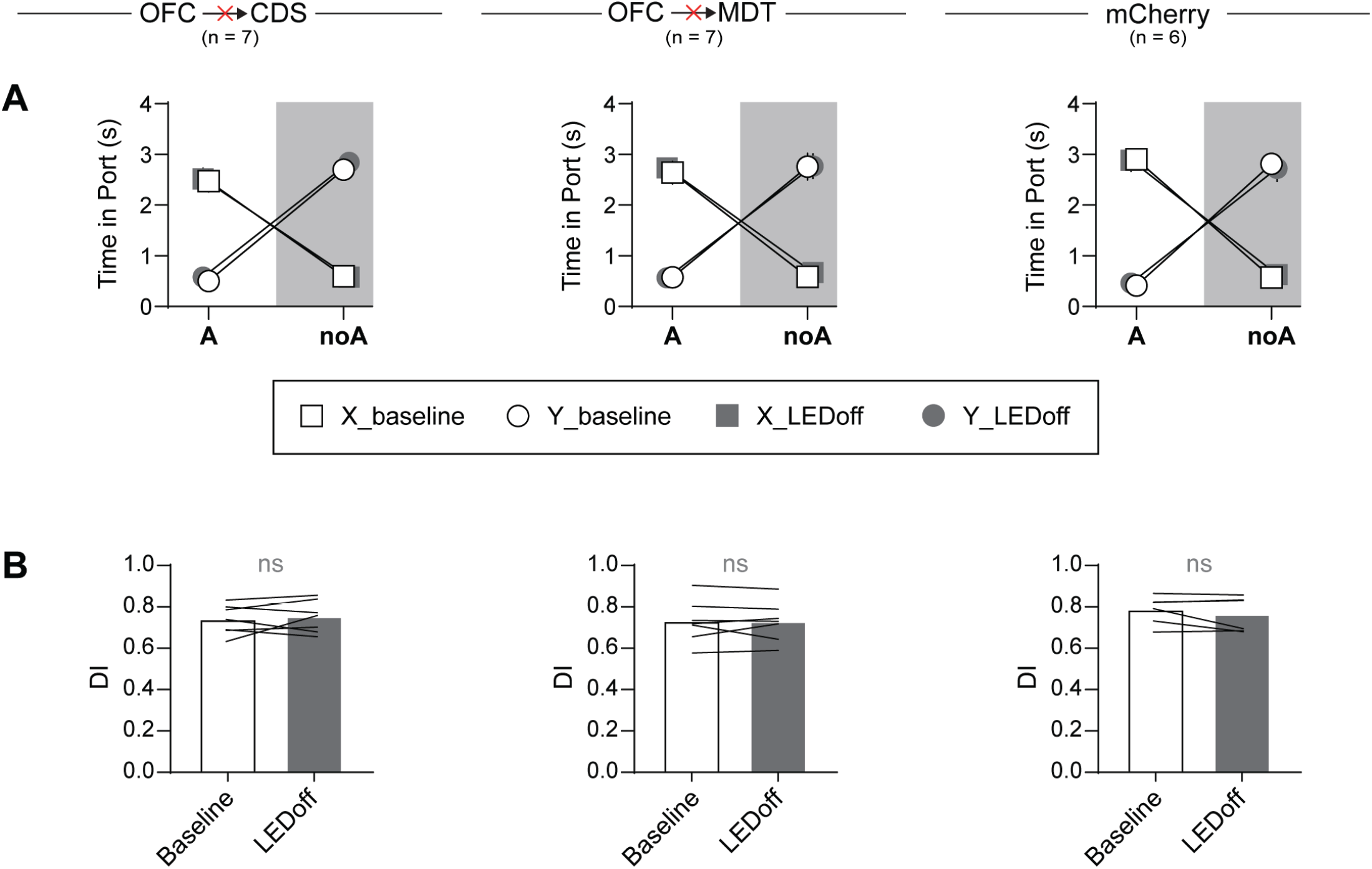
No evidence for carryover effects of phasic OFC output pathway inhibition beyond stimulated trials. **A**.Mean port occupancy during the first 5 s of cue presentation (prior to reward delivery) for each trial type (A:X, X, A:Y, Y), comparing LEDoff trials (within stimulation sessions) to baseline (no-stimulation) sessions. An omnibus linear mixed-effects model revealed no main effect of session type (baseline vs. stimulation session) and no interactions involving session type (Ps > 0.08). This absence of session-related effects was independently confirmed within each group using targeted linear mixed-effects models (Ps > 0.07). **B**.auROC-based discrimination index for baseline sessions and LEDoff trials. For each group, discrimination performance remained stable across sessions (Ps > 0.27; bootstrap tests). Collectively, these results rule out any lingering or carryover effects of LED stimulations, confirming that the effects of optogenetic manipulations were temporally specific and confined to stimulated trials.

**Fig. S3:**
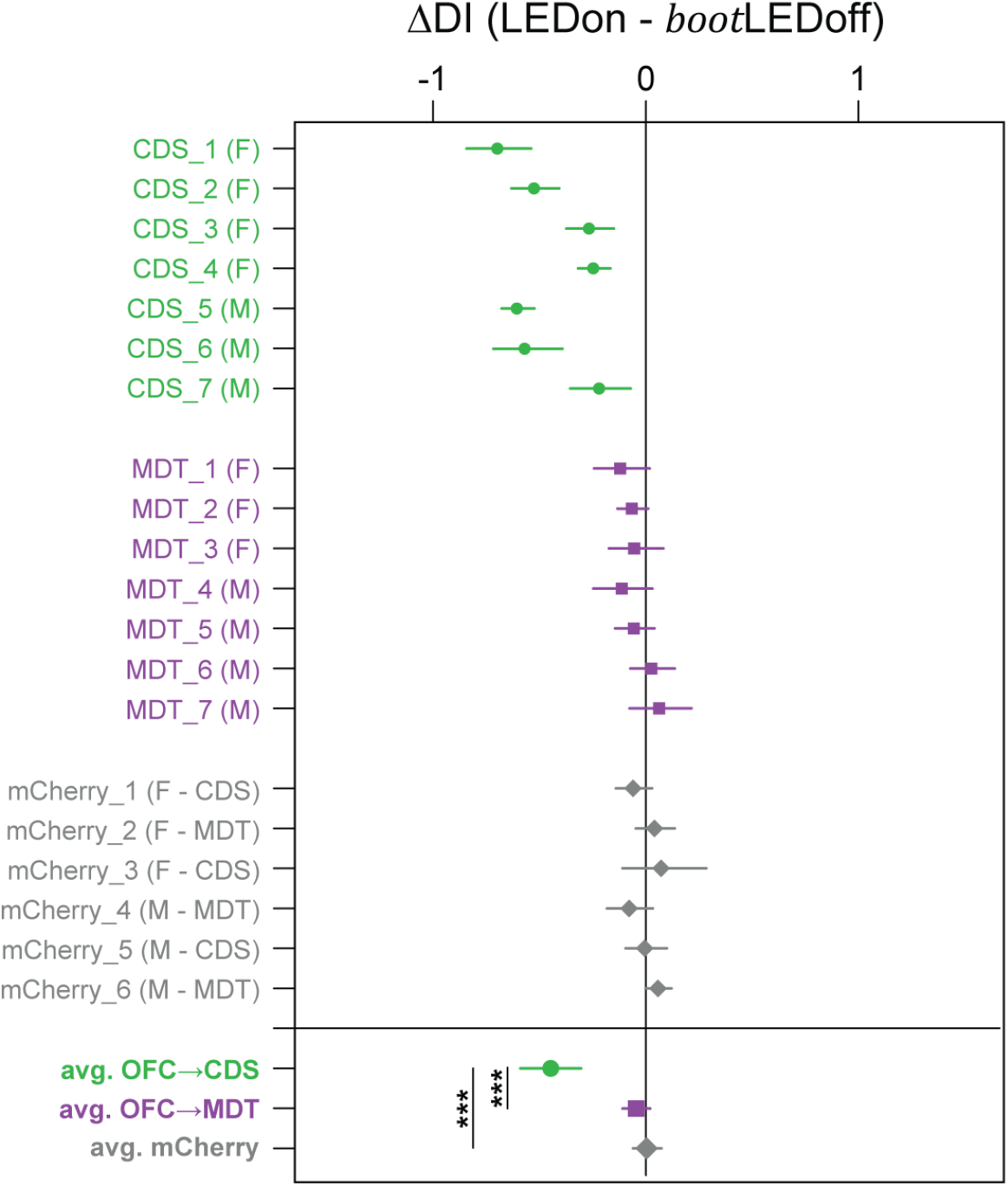
Summary effects of LED stimulation on discrimination index. Effects of LED stimulation on discrimination performance expressed as ΔDI (DI_LEDon – bootstrapped DI_LEDoff). Top: Subject-level estimates of ΔDI. Bottom: Group-level estimates of ΔDI. Symbols indicate mean values; error bars indicate 95% bootstrap confidence intervals. ***: P<0.001 bootstrap test.

**Fig. S4:**
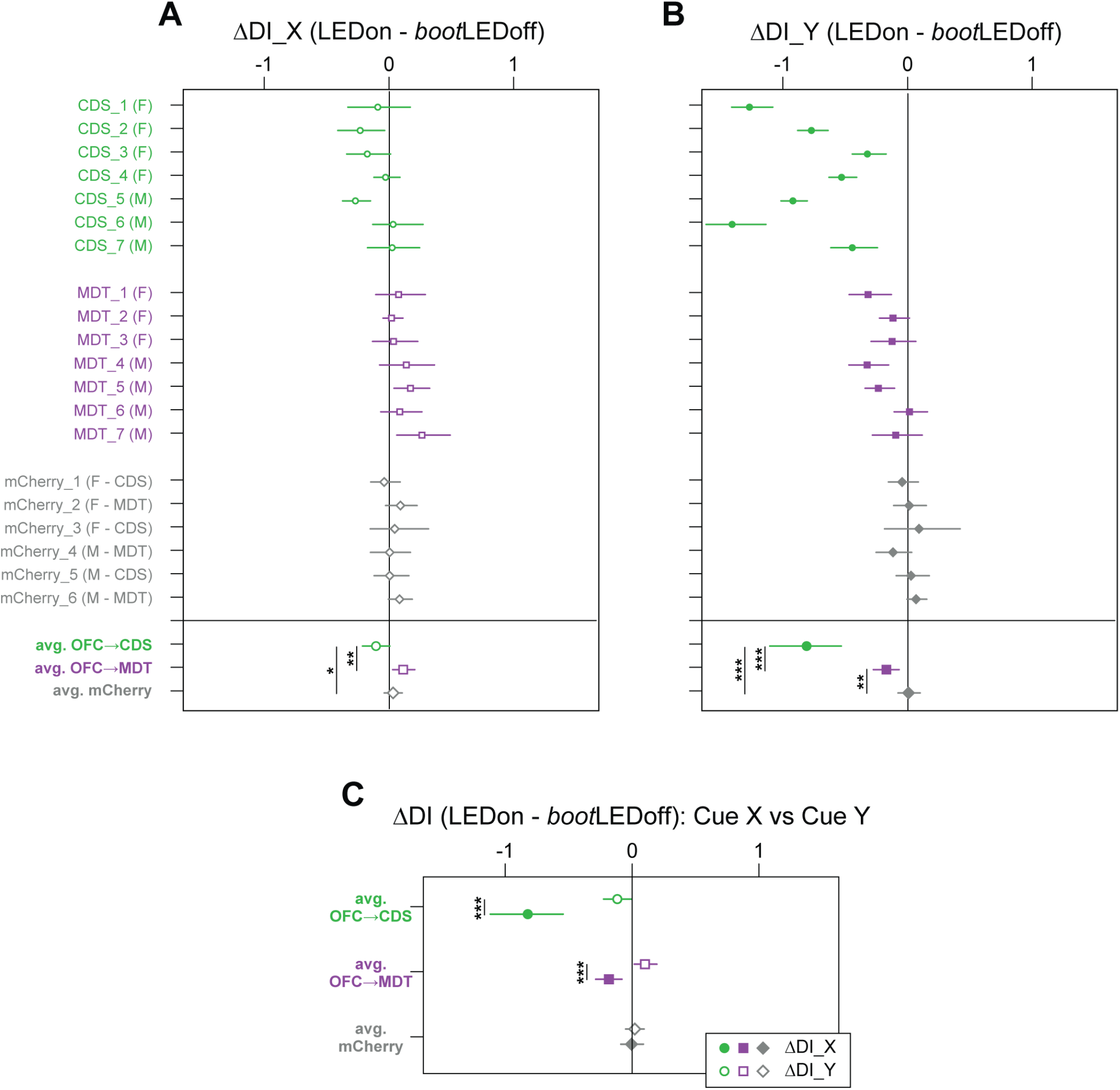
Summary effects of LED stimulation on discrimination index for each target cue. **A**.Effects of LED stimulation on discrimination performance for the A:X+ / X-component of the task. Top: Subject-level estimates of ΔDI. Bottom: Group-level estimates of ΔDI. **B**.Effects of LED stimulation on discrimination performance for the A:Y- / Y+ component of the task. Top: Subject-level estimates of ΔDI. Bottom: Group-level estimates of ΔDI. **C**.Direct comparison of LED stimulation effects on the two task components. Symbols indicate mean values; error bars represent 95% bootstrap confidence intervals. * P < 0.05; ** P < 0.01; *** P < 0.001; bootstrap test.

**Fig. S5:**
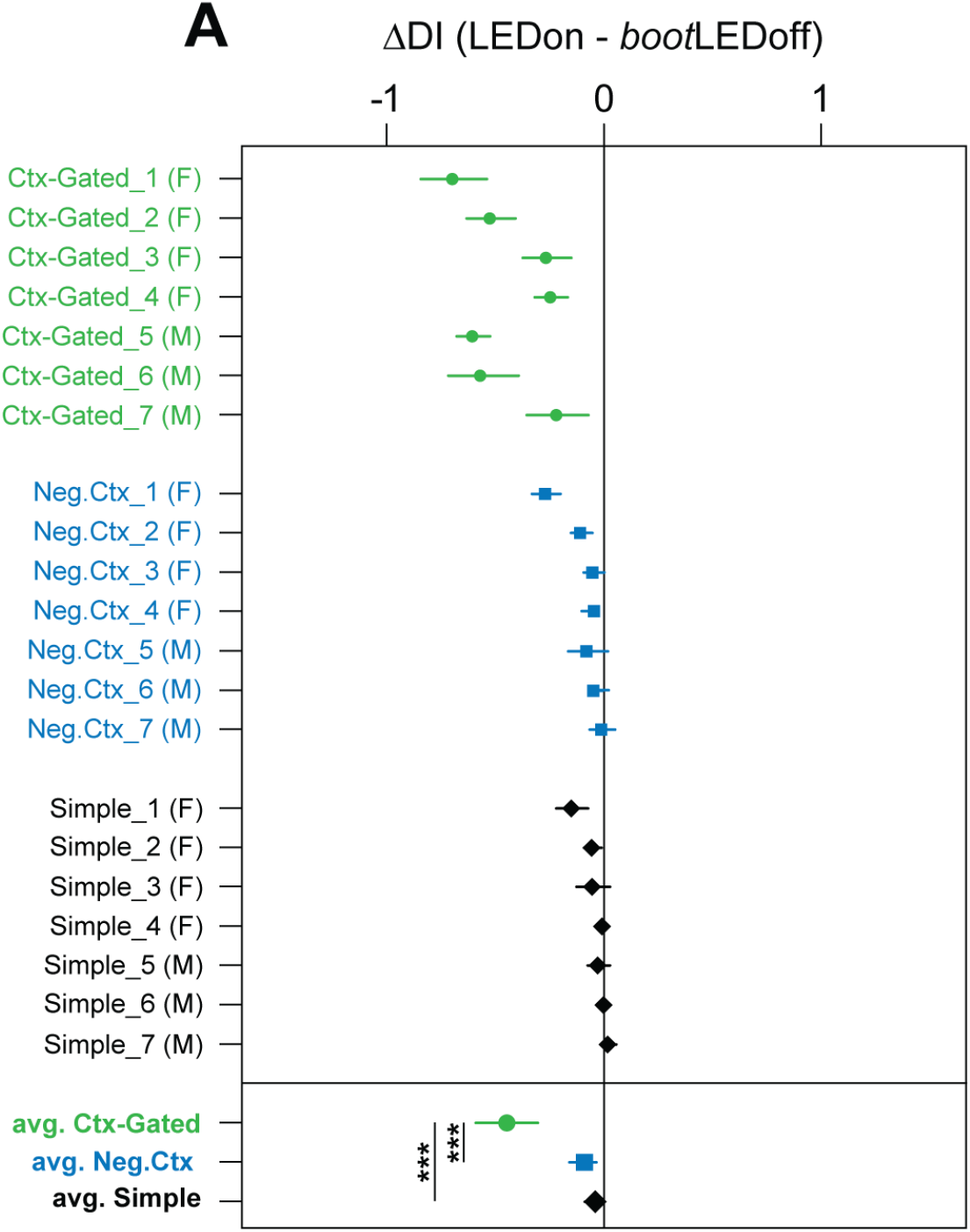
Summary effects of OFC→CDS silencing on discrimination index across behavioral task. Effects of LED stimulation on discrimination performance expressed as ΔDI (DI_LEDon – bootstrapped DI_LEDoff). Top: Subject-level estimates of ΔDI. Bottom: Group-level estimates of ΔDI. Symbols indicate mean values; error bars indicate 95% bootstrap confidence intervals. ***: P<0.001 bootstrap test.

**Fig. S6:**
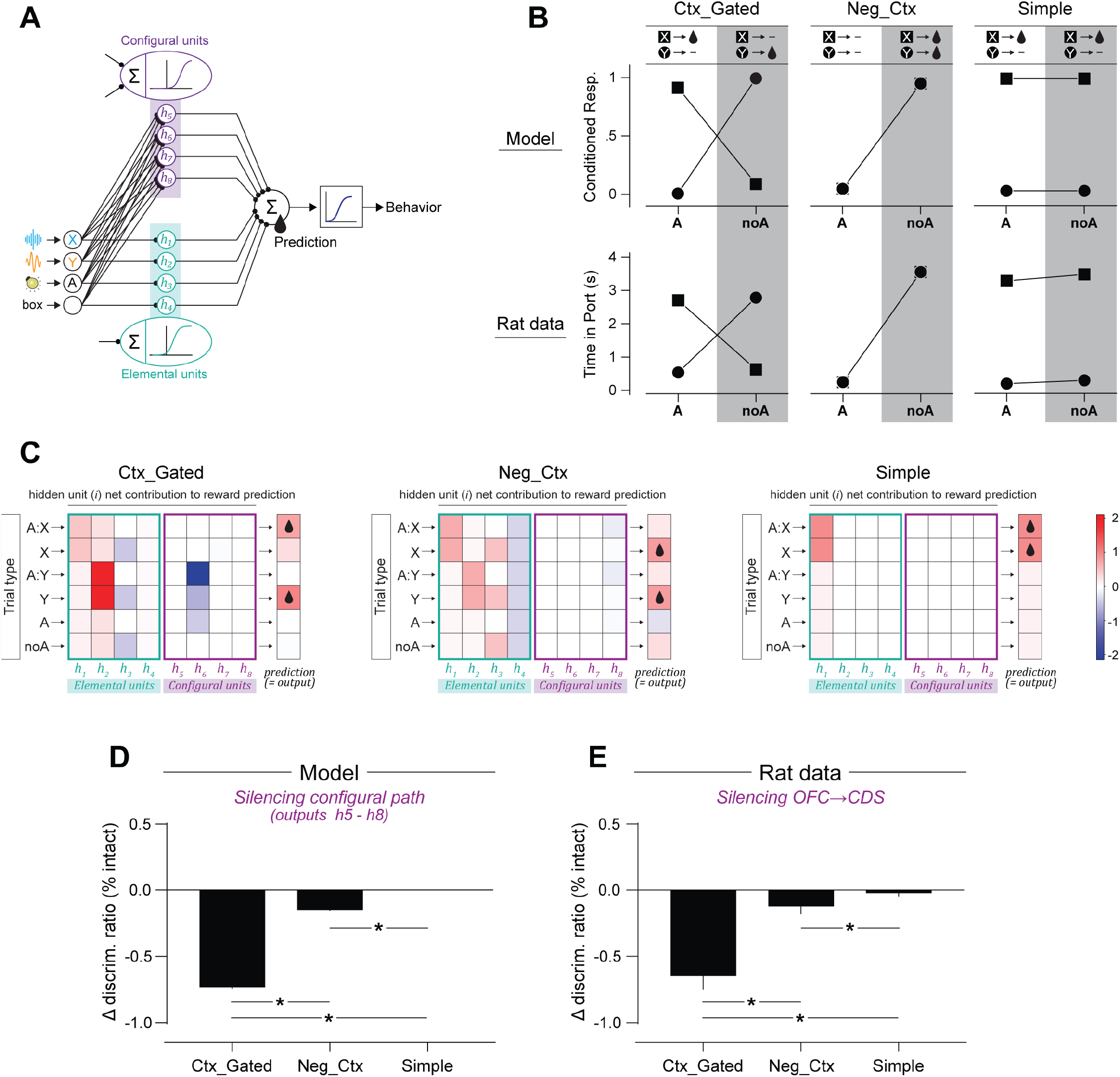
Connectionist model accounts for task-specific OFC→CDS contributions. **A**.Model architecture. **B**.Network performance (top) and rat behavior (bottom) after training to criterion in the three behavioral tasks: context-dependent discrimination (left); negative occasion-setting (middle); simple auditory discrimination (right). **C**.Hidden-unit contributions to outcome prediction. The contribution of each hidden unit (i) was computed as A_i_ × W_i_→out (A_i_: unit activity; W_i_→out: weight to output). Configural units are engaged primarily in the context-dependent discrimination task, with limited involvement in the negative occasion-setting task and virtually no contribution to the simple discrimination task. **D**.Simulated disruption of the configural pathway. Consistent with the task-dependent engagement of configural units, silencing the configural pathway strongly disrupted network discrimination in the context-dependent discrimination task, produced minimal disruption in the negative occasion-setting task, and had virtually no effect in the simple discrimination task. **E**. These simulated effects parallel the behavioral consequences of OFC→CDS silencing observed in rats across the same tasks.

**Table. S1:**
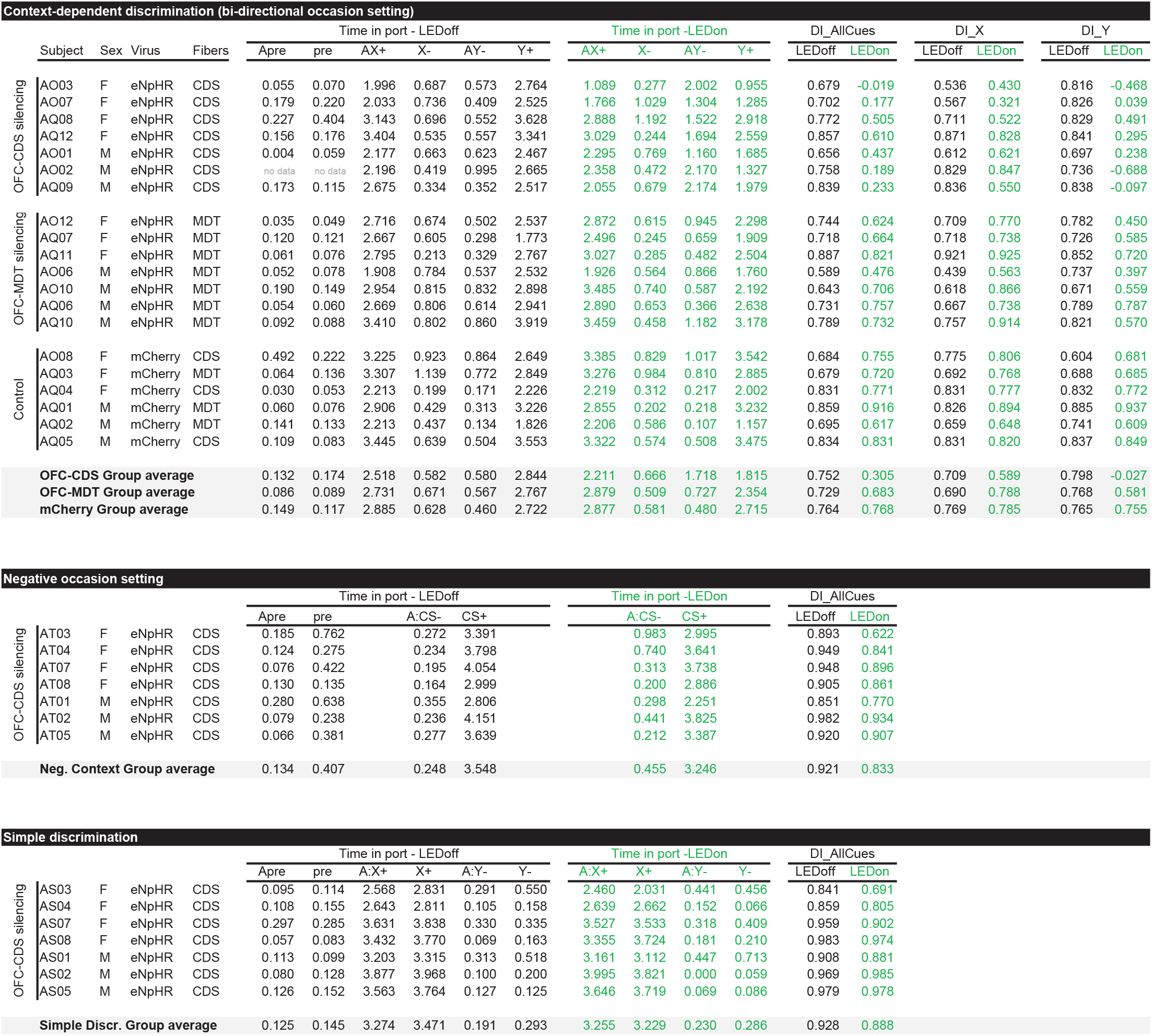
Subject-level mean port occupancy for each trial type.

## Notes

### Competing Interest Statement

The authors have declared no competing interest.

### Funding

- Hellman Foundation

